# Polyp bailout and reattachment of the abundant Caribbean octocoral *Eunicea flexuosa*

**DOI:** 10.1101/2020.07.04.187930

**Authors:** Christopher D. Wells, Kaitlyn J. Tonra

## Abstract

Anthozoans exhibit great plasticity in their responses to stressful conditions, including decreasing individual size, detaching from the substratum and relocating, and releasing endosymbiotic microalgae. Another response to stress used by some colonial anthozoans is polyp bailout, in which the coenenchyme breaks down and individual polyps detach from the colony. We observed polyp bailout in the common Caribbean gorgonian *Eunicea flexuosa* after eight hours of aerial exposure. After nine days, 28% of bailed-out polyps reattached, although none opened to resume feeding. Polyp bailout is a costly and high-risk escape response, but reattachment indicates that this can be a genet-saving behavior in cases where whole-colony mortality is likely. While it has been described in several species of scleractinians and two octocorals, we still do not know how widespread this behavior is in anthozoans.

## Introduction

As marine environments around the world change in response to warming climates and increasing severe weather events, coral reef organisms such as scleractinian corals and octocorals face stressful environmental conditions (Carpenter et al. 2008). Under thermal stress, corals respond by bleaching, becoming less resilient to disease, and slowing growth rates (Bruno et al. 2007; Cantin et al. 2010; Hughes et al. 2017). Another response to environmental stress in some coral polyps is through a temporary breakdown of their colonial structure when local conditions are too extreme. This is done by detaching from their colony mates and reattaching in a more suitable location in a process known as “polyp bailout” (*sensu* Sammarco 1982).

Polyp bailout is a directed breakdown of coloniality in some anthozoans through tissue-specific apoptosis (Kvitt et al. 2015). It can be triggered by starvation (e.g., Goreau and Goreau 1959), chemical stress (Yuyama et al. 2012), or large changes in salinity, temperature, pH, and oxygen levels (e.g., Kružić 2007; Kvitt et al. 2015; Shapiro et al. 2016; Rakka et al. 2019). Bailed-out polyps drift until finding a suitable attachment location and, if successful, resume colonial growth (Sammarco 1982). This behavior is different from “polyp expulsion” *(sensu* Kramarsky, 1997), a form of asexual budding where individual polyps and their calices detach from healthy colonies and then reattach to form new colonies.

Polyp bailout has been observed in seven scleractinian corals *(Acropora tenuis:* Yuyama et al. 2012; *Astroides calycularis:* Serrano et al. 2018; *Cladocora caespitosa:* Kružić 2007; *Pocillopora damicornis:* Kvitt *et al.* 2015; Shapiro et al. 2016; Fordyce et al. 2017; *Seriatopora hystrix:* Sammarco 1982; *Tubastrea coccinea:* Capel et al. 2014) and two octocorals *(Acanella arbuscula* and *Acanthogorgia armata:* Rakka et al. 2019). Reattachment has been observed in three of the scleractinians: *C. caespitosa* (Kružić 2007), *P. damicornis* (Shapiro et al. 2016), and *S. hystrix* (Sammarco 1982). Neither octocoral species reattached (Rakka et al. 2019). In this study, we observed polyp bailout after exposure to air for eight hours and subsequent reattachment of polyps of the abundant Caribbean octocoral *Eunicea flexuosa*. This description of polyp bailout is a contribution to the largely unexplored topic of stress responses in octocorals.

## Methods

Branches of *Eunicea flexuosa* were collected from an octocoral-dominated reef in Round Bay, St. John, U.S. Virgin Islands (18.345 °N, 64.681 °W) on 14-15 July, 2019 between 3.0 and 6.0 m depth. Colonies were maintained in a sea table (i.e., flow-through, open-topped aquarium) with unfiltered running seawater at the Virgin Islands Environmental Resource Station. For the first six weeks, colonies did not show signs of stress. Due to a sea table malfunction overnight, the water level was reduced so only the lowest five centimeters of the branches were submerged. The water level was returned to normal height after at most eight hours. Colonies were visibly stressed; polyps became fully extended and the coenenchyme changed from beige to dark brown. Within two days coenenchyme had begun to disintegrate. Five days after being exposed to air, bailed-out polyps were observed and collected with a 150-μm mesh filter attached to the outflow of the sea table. In order to determine if these polyps could reattach, 40 bailed-out polyps were placed in a 1-L dish with a stoneware clay tile (14 × 14 × 1 cm). The tile was examined for signs of attachment after nine days by gently jetting water from a transfer pipette at polyps that appeared to have reattached.

## Results and Discussion

Bailed-out polyps were slightly negatively buoyant, did not possess external sclerites, and were 550 to 650 μm wide. It is unknown whether polyps contained internal sclerites. They were an unusual grey-blue hue (Fig. 1). These observations of tissue contraction and loss of sclerites have been observed in two other octocorals where polyps detached after being under stress (Rakka et al. 2019). After nine days, 11 of the 40 polyps (28%) reattached to the tile (Fig. 2). There was no indication of axis redevelopment during the first nine days. The other 29 polyps remained unattached but had not died. Tentacles were never observed extended, indicating that the polyps were not feeding after bailing out of their colonies. Endosymbiotic dinoflagellates (Symbiodiniaceae) were visible (Fig. 1) and may have been providing a source of energy while the polyps were attaching to the substratum.

**Fig. 1.**
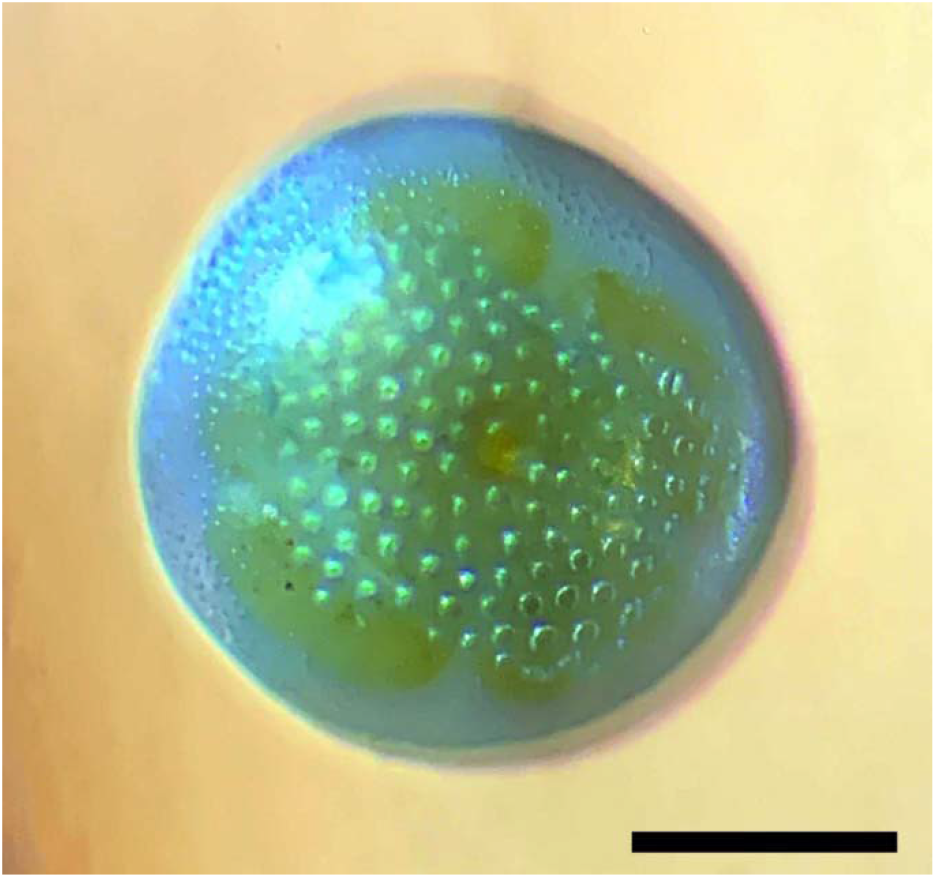
Aboral end of a bailed-out polyp of the gorgonian *Eunicea flexuosa* immediately after collection. Many clusters of golden-brown endosymbiotic Symbiodiniaceae are visible. Scale bar is 200 μm.

**Fig. 2.**
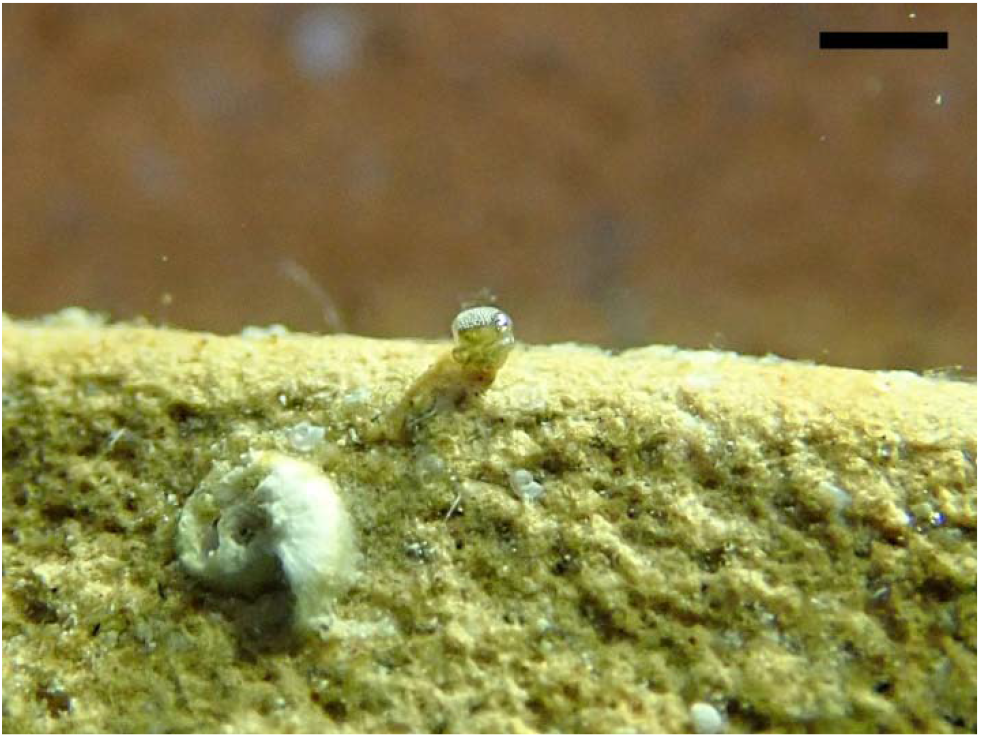
Reattached bailed-out polyp of the gorgonian *Eunicea flexuosa*. Scale bar is 1.0 mm.

Bailing out of a colony is a potential alternative to death, but there are many inherent risks with this strategy and the loss of coloniality. Octocorals generally follow a type III survival curve (Linares et al. 2007), meaning that mortality risk is much higher for single polyps than adult colonies. Additionally, most octocorals have to reach a minimum size before being able to reproduce (Kahng et al. 2011). In scleractinians, damage or asexual reproduction that results in a reproductive colony becoming smaller than this minimum size can lead to the colony becoming non-reproductive (i.e., “reverse puberty”) until it surpasses the minimum reproductive size (Szmant 1986). Despite these costs, several anthozoans have the capability of disassociating from their colony during stressful times (e.g., Capel et al. 2014; Kvitt et al. 2015; Shapiro et al. 2016) and it may be a normal form of asexual reproduction for some corals (Sammarco 1982). While there are other forms of asexual reproduction in octocorals, such as autotomizing of branches (Lasker et al. 1984), we do not believe that *E. flexuosa* is using this behavior as a method of reproduction, but rather as a genet-saving behavior. It seems that coenenchyme degradation, often seen in diseased and damaged octocorals (e.g., Morse et al. 1981; Wahle 1985; Harvell and Suchanek 1987), and subsequent polyp bailout, is a potentially viable last-ditch option for octocorals to survive an otherwise fatally stressful experience.

Currently, we do not know how widespread this behavior is in octocorals or whether this behavior is simply an artefact of laboratory culturing. Studies that induce polyp bailout in scleractinians have already shed light on important phenomena such as calcification, bleaching, and disease in corals (Shapiro et al. 2016), all critical topics in the face of the current coral reef crisis (Bellwood et al. 2004; Veron et al. 2009; Hughes et al. 2010). Indeed, a more widespread survey of this behavior is needed within Octocorallia, especially *in situ* during stressful events such as reef-wide bleaching. With the potential change of scleractinian coral dominated reefs to octocoral dominated reefs in the Caribbean (e.g., Ruzicka et al. 2013; Lenz et al. 2015; Edmunds and Lasker 2016; Sánchez et al. 2019), it is critical to understand the physiological impacts of environmental stress on octocorals.

## Acknowledgements

This work was completed under permits from the Virgin Islands National Park (VIIS-2019-SCI-0011) and the Virgin Islands Division of Fish and Wildlife (DFW19010) and was funded by the National Science Foundation (OCE 1756381). We thank the Virgin Islands Environmental Research Station and the University of the Virgin Islands for laboratory space, L. Bramanti, H.R. Lasker, Á. Martínez-Quintana, and A.M. Wilson for field and laboratory assistance, and E.R. Anderson, H.R. Lasker, and Á. Martínez-Quintana for reading and commenting on the manuscript. The authors declare that there is no conflict of interest regarding the publication of this article.

